# Antibiotic Resistance in *Staphylococcus aureus*: Effects of Quorum Sensing Inhibition and DNA Fragmentation

**DOI:** 10.1101/2025.04.26.649666

**Authors:** Sreya Kurup, Patrick K. Taylor, Paul J. Adams, Barnabe D. Assogba

## Abstract

**Background:** Antimicrobial resistance (AMR) is a global crisis, causing 2.8 million infections and 35,000 deaths annually. Staphylococcus aureus is mainly responsible for causing these challenging infections through biofilm formation and the action of efflux pumps. A limited number of studies on Hop (Humulus lupulus) have shown its potential to inhibit quorum sensing in pathogenic bacteria.

**Objective:** Therefore, a novel treatment approach was used in this study, which investigated Hop’s β-acids, particularly the combination of colupulone and n+adlupulone, as well as in combination with fluoroquinolone antibiotics ciprofloxacin and ofloxacin. As ciprofloxacin remains a highly effective antibiotic against Staphylococcus aureus but resistance can develop, and ofloxacin exhibits naturally higher resistance in S. aureus, this study hypothesized that combining Hop (containing colupulone & n+adlupulone) with the two antibiotics separately would result in a greater reduction in biofilm growth of S. aureus compared to their individual potency levels.

**Methods:** Antimicrobial activity was assessed using disk diffusion assays and minimum inhibitory concentration for biofilms at multiple concentrations through 2-fold serial dilutions.

**Results:** Our data demonstrate that Hop-derived β-acids possess direct antimicrobial activity and when combined with the fluoroquinolone antibiotics, exhibit additive or synergistic effects by acting on different targets in Staphylococcus aureus.

**Conclusions:** This study provides insight into how natural products can potentially mitigate the development of resistance to antibiotics like ciprofloxacin in the highly pathogenic bacterium S. aureus. It also highlights how adding natural compounds could improve drug effectiveness. Therefore, this demonstrates the potential of natural compounds and antibiotics like ofloxacin, which are known to be ineffective against S. aureus. It offers a promising natural-conventional hybrid approach to addressing antimicrobial resistance.

## Introduction

Antimicrobial resistance (AMR) is a global crisis, causing 2.8 million infections and 35,000 deaths annually [1]. Driven by antibiotic overuse, AMR complicates treating common infections and performing surgeries as the pathogen increasingly resists treatment. Particularly, bacterial AMR caused 1.27 million global deaths in 2019 alone, which previous studies described as a “Silent Pandemic” created by the misuse of antibiotics in humans, animals, and in agriculture, leading to the spread of antimicrobial-resistant genes and pathogenic bacteria [2,3]. Additionally, the World Health Organization (WHO) has identified antimicrobial resistance as one of the top three threats to human health, highlighting its impact [4].

*Staphylococcal species* are responsible for ∼60% to 70% of challenging AMR infections, of which *S. aureus* is a significant contributor [5]. S. aureus, a gram-positive pathogen, has developed significant antibiotic resistance, particularly fluoroquinolones, through mechanisms such as biofilm formation and efflux pump activity [6].

In addition, the *blaZ* gene, encoding a beta lactamase, is a significant contributor for penicillin resistance and development of highly drug resistant “superbug” strains of methicillin-resistant *S. aureus* (MRSA) [7]. *S. aureus* evades the host immune system by forming biofilms, through bacterial communication systems that regulate the population density by autoinducers, called Quorum Sensing (QS). After a threshold of growth, QS activates genes, causing the production of extracellular polymeric substances (EPS) in bacteria, providing support and protection to the biofilm, leading to the bacteria adhering to surfaces [8]. Biofilms can hinder antibiotic effectiveness by creating a protective barrier where cells in the outer layers, with higher metabolic activity, are exposed to the antibiotic. These cells deplete oxygen and nutrients, leaving the deeper cells with decreased antibiotic exposure and limited metabolic activity. As the outer cells are killed off, the deeper cells known as persisters, rapidly regrow, making infections harder to treat and potentially more resistant than before. This can lead to reduced local concentrations of antimicrobial compounds and evasion of the immune system which results in incomplete clearance of an infection. Unlike other bacteria, *S. aureus* can cause serious infections if it invades deeper into the body through your bloodstream [9]. These biofilms result in major infections like osteomyelitis, endocarditis, and device-related infections, while complicating treatments, resulting in longer hospital stays and higher mortality rates [10].

Hop (*Humulus lupulus*), a natural plant commonly used for 98% of the beer brewing industry worldwide, mainly uses the female cones of Hop called the strobili. Hops were introduced during the seventeenth century in North America– currently, the United States is the biggest producer of Hop, contributing to more than half of the Hop produced annually, worldwide [11]. Hops were traditionally used as sedatives and currently they have been approved as treatment for restlessness, anxiety, and sleep disturbances [12]. Recent studies are now noticing the effectiveness of Hop’s compounds and exploring it against oral health and cancer. Previous studies have shown Hop’s natural antibiofilm activity, through triggering the quorum-sensing-dependent gene expression by binding to its signal peptides of the cell membrane, therefore potentially resulting in quorum-sensing inhibition [13].

Fluoroquinolones are characterized into four different generations, where different fluoroquinolones inhibit a spectrum of different antibacterial activities [14,15]. Before the introduction of newer fluoroquinolones, the potency of fluoroquinolones against gram-positive bacteria was inconsistent, after which ciprofloxacin and ofloxacin were introduced which was significantly effective against *S. aureus*. Both ciprofloxacin and ofloxacin hinder the production of pathogens by targeting DNA gyrase and topoisomerase IV resulting in DNA fragmentation. Ciprofloxacin and ofloxacin are also heavily used because of their range of activity and great oral bioavailability [16,17]. Although the overuse of the antibiotics have led to mutations resulting in reduced drug accumulation through increased efflux, causing antibiotic resistance. This resistance reduced the effectiveness of both ciprofloxacin and ofloxacin in treating infections caused by resistant bacteria, such as *S. aureus* [18].

Combination therapy further enhances the potency of a single drug by targeting multiple pathways, therefore reducing the resistance. This has been proven to be effective against bacterial infections such as *Mycobacterium tuberculosis*, for which four-drug combinations are used. Currently, monotherapy fails to provide protection against resistant infections because of the rapid mutations and adaptations of the resistant pathogens, resulting in patient treatment complications. This failure of monotherapy, especially in *S. aureus* is largely because of biofilms and the efflux pump mechanism, which plays a crucial role in bacterial AMR. This mechanism was observed very early on in *S. aureus* and *Streptococcus pneumoniae* against ciprofloxacin, which is known to be one of the most effective antibiotics, especially compared with the potency of ofloxacin against *S. aureus* [19].

The limited effectiveness of conventional antibiotics such as ciprofloxacin and ofloxacin against resistant bacteria like *S. aureus* is the gap this study aims to bridge by investigating the effects of Hop—particularly the mixture of colupulone and n+adlupulone compounds—in combination with ciprofloxacin and ofloxacin [20]. Therefore, this study targets both the sessile and planktonic forms, rather than solely relying on individual treatments. It was hypothesized that the combination treatment would provide a higher inhibition against both the sessile and planktonic forms of *S. aureus*, by causing quorum sensing inhibition and inducing DNA fragmentation, compared to the potency levels of just the antibiotics alone. This perspective is supported by recent studies, which highlight that combination therapies can potentially improve treatment outcomes and reduce resistance, promoting the need for innovative strategies in managing resistant infections [2].

## Materials and Methods

### Bacterial strain & suspension preparation

The *S. aureus* ATCC 25923 strain was quadrant streaked and incubated overnight at 37°C to isolate single colonies, which were inoculated for preparing bacterial suspensions in 4 ml of Tryptic Soy Broth (TSB) according to laboratory guidelines. This was then further incubated at 225 RPM, and bacterial suspensions were normalized by taking the optical density at 600 nm (OD600) for both disk diffusion and minimum inhibitory concentration (MIC) assays. Biofilm formation was assessed using crystal violet staining after treatment with serial concentrations of compounds for the MIC assay.

### Extraction, dilution & identification of Hop compounds

The commercial product Hop Guard 3 (Manufactured by Dancing Bee Equipment), a viscous mixture of Hop extract, was used as a source of *β-acids* in this study. Subsequently, this viscous Hop extract was measured and vortexed with methanol, resulting in a homogenous liquid extract, with the total concentration being 2500 µg/ml for disk diffusion assays, and 2048 µg/ml concentration for the biofilm assay. In addition, High-Performance Liquid Chromatography (HPLC) was performed to identify whether the hop extract only consisted of colupulone and n+adlupulone compounds, as required for this study. As observed in (Figure 1), the HPLC showed that the compounds present in the Hop extract consisted only of colupulone and n-adlupulone. Therefore, this extract was further used in the concentrations of the respective assays. Herein, this Hop extract in methanol is referred to as Hop in experimental conditions.

**Figure 1:**
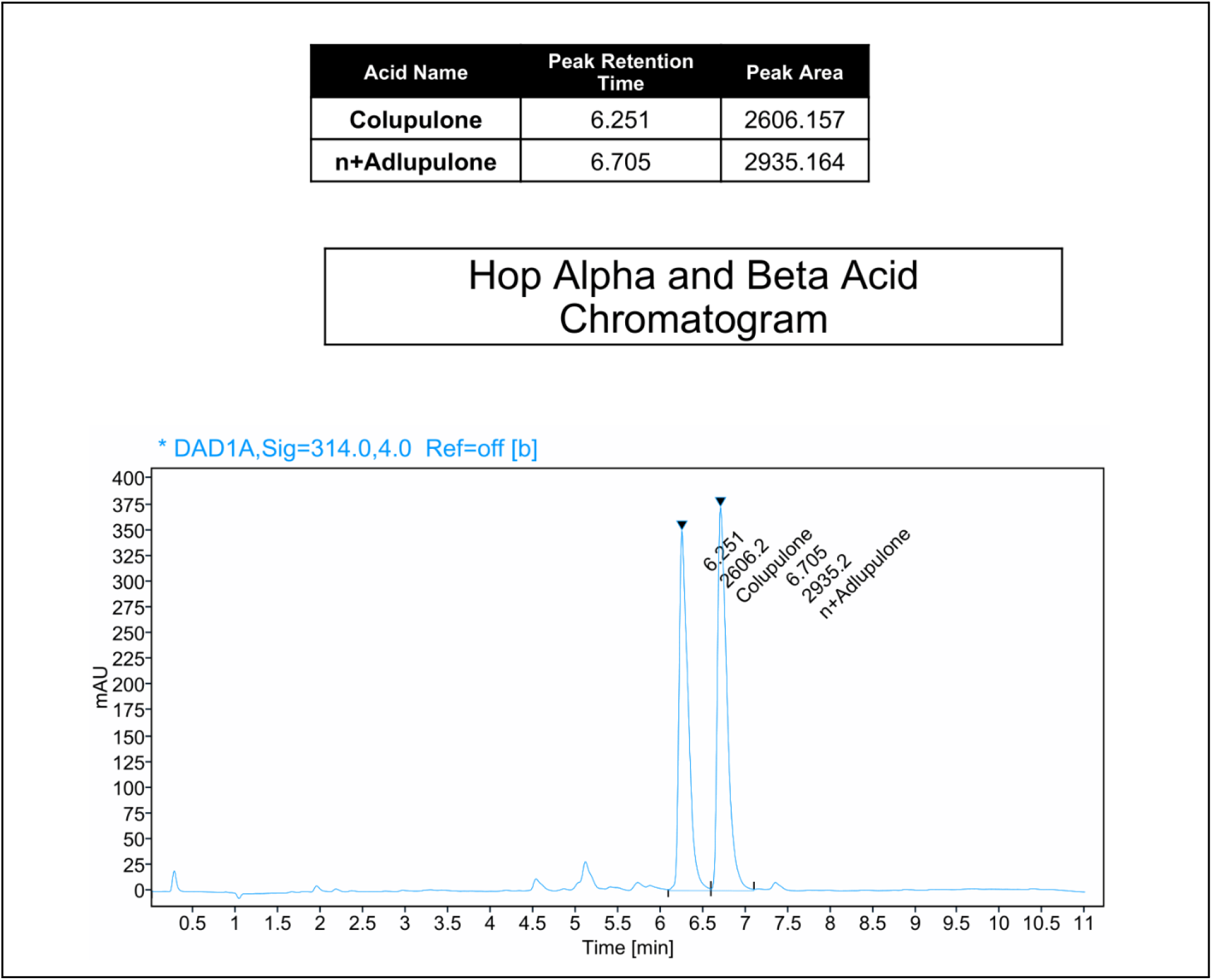
HPLC results of the Hop extract, suggesting only the presence of Colupulone and n+Adlupulone, as required for this study.

### Disk diffusion assay

Disk diffusion was performed as described by Jan Hudzicki with modifications [21]. Seven different treatments (1 disk/plate) were tested against S. aureus sessile form, where bacteria adhered to a surface but were not yet in a biofilm form. Each treatment was repeated eight times to ensure statistical accuracy, as seen in Figure 2. The seven treatment groups included methanol and sterile ultrapure distilled water as the negative control group; Hop, ciprofloxacin and ofloxacin as the individual comparison groups; alongside Hop with ciprofloxacin and Hop with ofloxacin as the combination groups. Rather than using pre-prepared disks, this study pipetted 20 µL of the respective treatment into each disk. Each plate contained 100 µL of the bacterial suspension, which was homogeneously spread using a cell spreader on the TSB agar plate before adding the disk. The concentration of the antibiotics used was determined according to the European Committee on Antimicrobial Susceptibility Testing (EUCAST), where the suggested concentrations of both Ciprofloxacin and Ofloxacin are 5 µg/10µL (solvent: pure water), against *S. aureus* [22]. Therefore, each disk of antibiotics consisted of 500 µg/ml, while the concentration of Hop was adapted from two different studies. Agnieszka Bartmańska’s study explored Hop’s antimicrobial effectiveness at lower concentrations of 50–100 µg, and a study by Di Lodovico *et al* (2020) investigated the concentration of Hop up to 5 mg/ml in disk diffusion assays, both being effective against *S. aureus* [23,24]. This study, therefore, explored the middle-range concentration of 2500 µg/ml, where 10 µL of methanol was added to 25 µg of Hop extract. Finally, to test the combination treatments, 10 µL of 500 µg/ml (respective antibiotic) and 10 µL of 2500 µg/ml (Hop) were pipetted into each disk and incubated for 24 h. Afterwards, ImageJ measured the zone of inhibition and checked for accuracy. Finally, the petri dishes were again incubated for five days to obtain data on how the treatments would react over an extended time against *S. aureus*.

**Figure 2:**
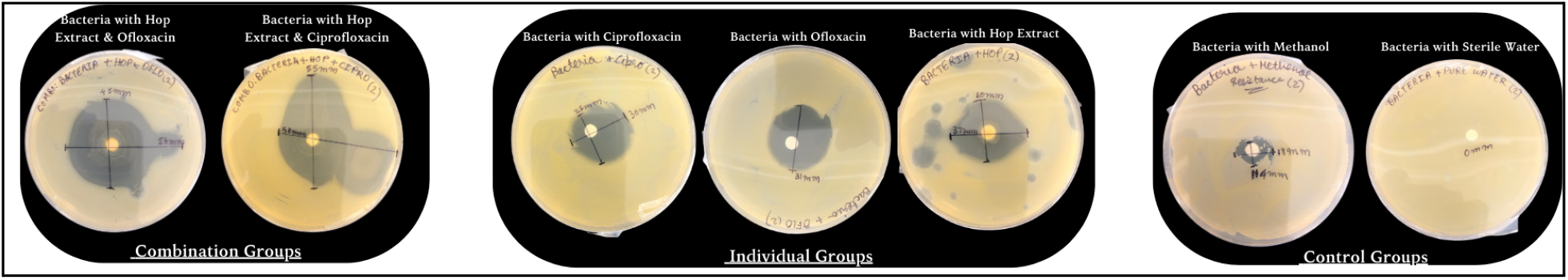
Seven treatment groups were tested against *S. aureus’s* sessile form, consisting of Hop+ofloxacin and Hop+iprofloxacin as the main combination groups; just ciprofloxacin, ofloxacin, and Hop as the individual comparison groups; and methanol and sterile water as the negative control groups. As seen in this figure, the largest zone of inhibition is present in the combination groups, representing the higher potency levels against *S. aureus* compared to just the individual treatments. Each treatment was repeated eight times to ensure statistical accuracy.

### 2-Fold serial dilution for microtiter plate biofilm assay

A 2-fold serial dilution process was followed for the MIC biofilm assay, where the highest concentration of Hop, antibiotics and the combination groups were prepared. The highest concentration used for Hop was 2048 µg/ml, which was determined by the findings of a study by Bogdanova *et al* (2018). In this assay, planktonic bacteria (suspended in liquid culture) were used to form biofilms, allowing for the evaluation of antimicrobial effects on both the suspended form (planktonic) and the biofilm-forming capacity of S. aureus. The biofilm formation was assessed using serial concentrations of the treatments in the microtiter plates. However, in Bogdanova *et al*.*’s (2018) study, the most significant inhibition was seen at 125 μg/ml of Hop (lupulone), which was adapted as 128 μg/ml* into this study. This was done to provide the middle concentration for 2-fold serial dilutions ranging from 2048 to 2 µg/ml [13]. For ciprofloxacin and ofloxacin, the highest starting concentration was 1.6 µg/ml, which was determined according to EUCAST guidelines. Two-fold dilutions of antibiotics were made from 1.6 µg/ml to 0.001 µg/ml. As for the combination groups, the concentrations of both the Hop and the antibiotics were mixed, for example, the highest starting concentration for the combination group was 2048 µg/ml (Hop) + 1.6 µg/ml (ciprofloxacin/ofloxacin), down to 2 µg/ml + 0.001 µg/ml. All concentrations were prepared by diluting the stock solution of antibiotics and Hop with TSB (including 1% glucose) as the solvent rather than diluting it in water or methanol, unlike disk diffusion.

### Crystal violet microtiter plate MIC biofilm assay

MIC biofilm assay was performed according to the Wiegand et al (2008) protocol with some modifications [25]. The treatment groups tested against the *S. aureus* biofilms were the combination and individual groups, along with a control group where only the biofilm was allowed to grow without any treatments. The 96-well plates were divided into two horizontal parts, where each plate consisted of two treatments, and the last 12th column was used as the control group. Except for the first column, the rest of the wells were allotted 100 μl of TSB with 1% glucose to ensure strong biofilm growth. Then 200 μl per well is aliquoted to the first column, according to the respective treatments, which also contained TSB and glucose, as mentioned above. Two-fold serial dilutions were performed using a multi-channel pipette, and 100 μl was transferred from the first column to the second column, and so on till the 11th column, after which the 100 μl of treatment was discarded for the 12th column to maintain a No Treatment control group. Pipetting up and down in each well before transfer to the next was done to ensure a homogeneous mixture. Finally, 1 μl of liquid *S. aureus* culture was inoculated into all the wells. This resulted in a total volume of 101 μl in each well, which was incubated at 37°C for 48 h for the biofilms to establish. After the incubation period, the plates were stained using 0.5% crystal violet (CV), for which the methodology was adapted from O’Toole (2011) with some modifications [26]. Firstly, the incubated cultures are removed from 96-well plates, using a multi-channel micropipettor; thereafter, these plates are washed three times with 100 μl of deionized water per well, after which 110 μl of 0.5% CV is added and the plate is left to stain for 15 minutes. CV is removed, and the plate is washed with deionized water to remove excess stain. Finally, 110 μl of 70% ethanol is added to each well to solubilize the CV staining of the biofilms. This was then quantified on a multi-mode plate reader using 595 nm wavelengths and used for statistical quantifications. This process was repeated for each concentration in the 2-fold serial dilution, ensuring 12 replicates, including biological and technical replicates.

## Results

### The highest zone of inhibition after 24 hours, against sessile S. aureus among the treatment groups

After 24 hours from the disk diffusion assay, the results showed that the combination treatments caused a greater zone of inhibition against S. aureus than their respective individual antibiotic treatments (Figure 3A). The highest median zone of inhibition can be seen in the Hop and ciprofloxacin combination treatment (54 mm), followed by the combination treatment of Hop with ofloxacin (46 mm). While the individual treatments provided comparatively lower amounts of inhibition, with Hop having the highest (32 mm), followed by ciprofloxacin (25 mm) and ofloxacin (26 mm), which demonstrated almost the same amount of inhibition. The negative control groups, water (no inhibition) and methanol (11 mm), provided the least amount of inhibition, therefore providing a standpoint comparison between the rest of the treatments. A one-way ANOVA was performed to ensure statistical significance of the results obtained, and a *p*-value < 0.05 between groups was found. Thereafter, to pinpoint which particular groups were significant, a Tukey’s post-hoc test was conducted, and the results showed that the combination groups were critical to both the individual antibiotic treatments and the negative control groups (shown in Figure 3A by an “asterisk”).

**Figure 3:**
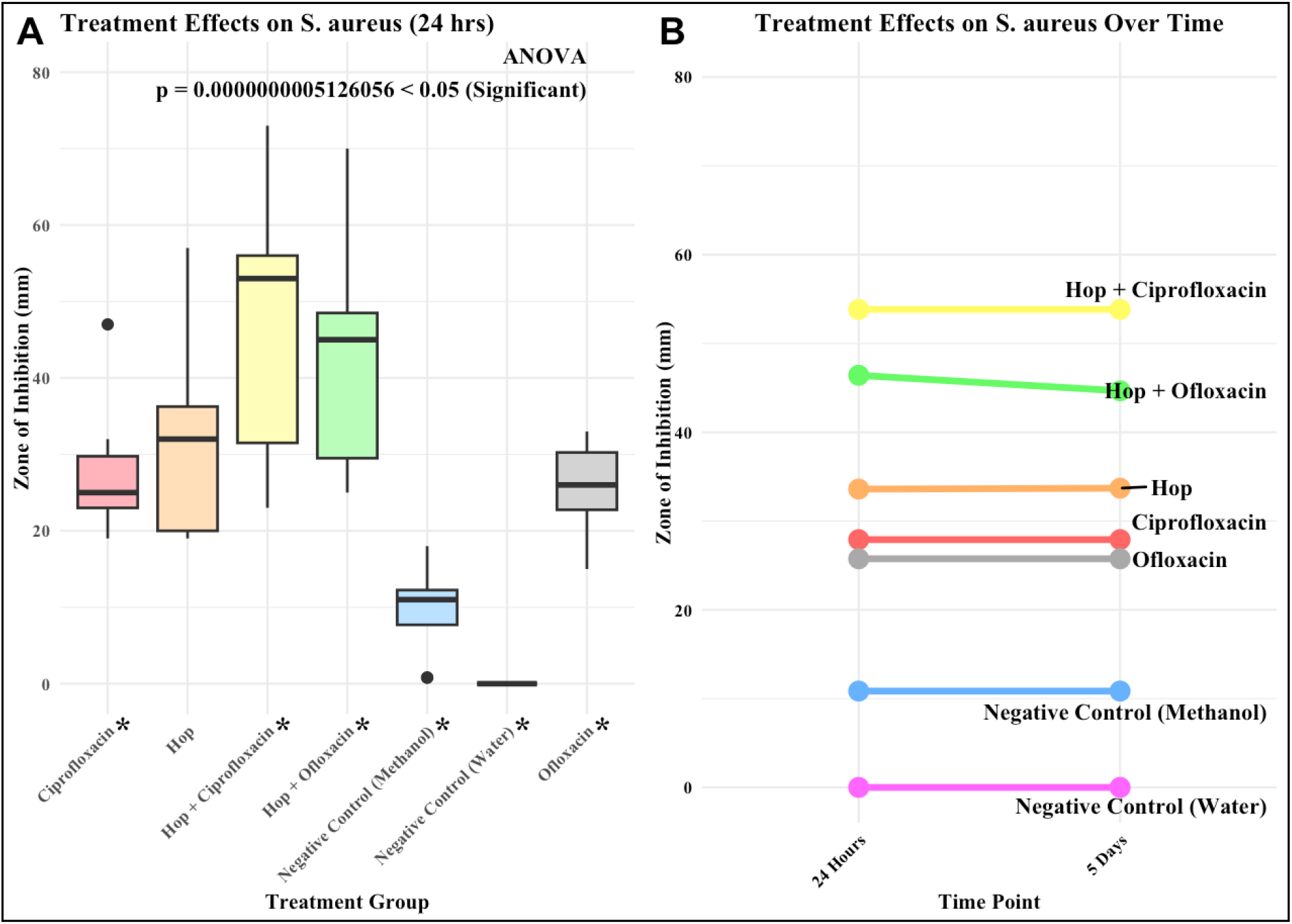
(A) The box plot shows the quantitative results obtained after 24 h of measuring the zone of inhibition of all seven treatment groups against the sessile *S. aureus*. The highest inhibition was observed in the hop combination with ciprofloxacin (54 mm), followed by the hop with ofloxacin (46 mm). Individual treatment groups of just hop (32 mm), ciprofloxacin (25 mm), ofloxacin (26 mm), and finally, the negative control groups had the least amount of inhibition. A one-way ANOVA was performed, which provided a *p*-value < 0.05. Individual treatment comparisons were tested with Tukey’s post-hoc, *p*-value < 0.05 shown by an “asterisk”. (B) The change in inhibition zones over time from 24 h to 120 h (5 days). Methanol and water acted as negative treatment controls.

### Changes to the zone of inhibition after five days (120 hours)—against sessile S. aureus

Surprisingly, the data obtained from the disk diffusion assay after five days (120 h) showed no drastic decreases in the potency levels of the seven treatment groups. As observed in Figure 3B, only slight changes were observed in all treatment groups. Combination groups displayed the highest potency, with Hop+ciprofloxacin changing from 54 mm to 53.8mm, followed by Hop+ofloxacin changing from 46 mm to 44.7 mm. The individual groups did not exhibit any significant changes; both Hop and ciprofloxacin remained the same (32 mm and 25 mm), followed by ofloxacin changing from 26 mm to 25.8 mm. As for the negative control groups, methanol changed from 11 mm to 10.8 mm, and pure water demonstrated no observable inhibition throughout. Among the combination treatments, the Hop+ofloxacin showed less sustainability, while Hop+ciprofloxacin’s combination inhibited the growth of *S. aureus* the most, sustaining it over 5 days of incubation.

### Highest biofilm inhibition after 48 hours, against planktonic S. aureus among treatment groups

The results obtained after 48 h from the MIC biofilm assay showed that the combination treatments significantly inhibited *S. aureus* biofilm, compared to the individual groups (Figure 4). The no-drug column (Figure 4B) demonstrated dense biofilm growth, represented by the dark purple colour, relative to individual antibiotic treatments (Figure 4D). Ofloxacin demonstrated a low level of inhibition relative to ciprofloxacin for all S. aureus biofilms, with the lightest shade observed at 0.05μg/ml, ciprofloxacin inhibited minimally at 0.001 μg/ml and 0.006 μg/ml to 0.0125 μg/ml, causing the majority of results to have dense biofilm growth still. Subsequently, Hop’s potency levels show an inhibition at 96μg/ml to 768μg/ml, as seen in Figure 4C. This range of activity with Hops resulted in a window of inhibition. Comparing all individual treatments against the combination treatments, it was observed that cultures treated with combinations of antimicrobials had considerably less *S. aureus* biofilm growth.

**Figure 4:**
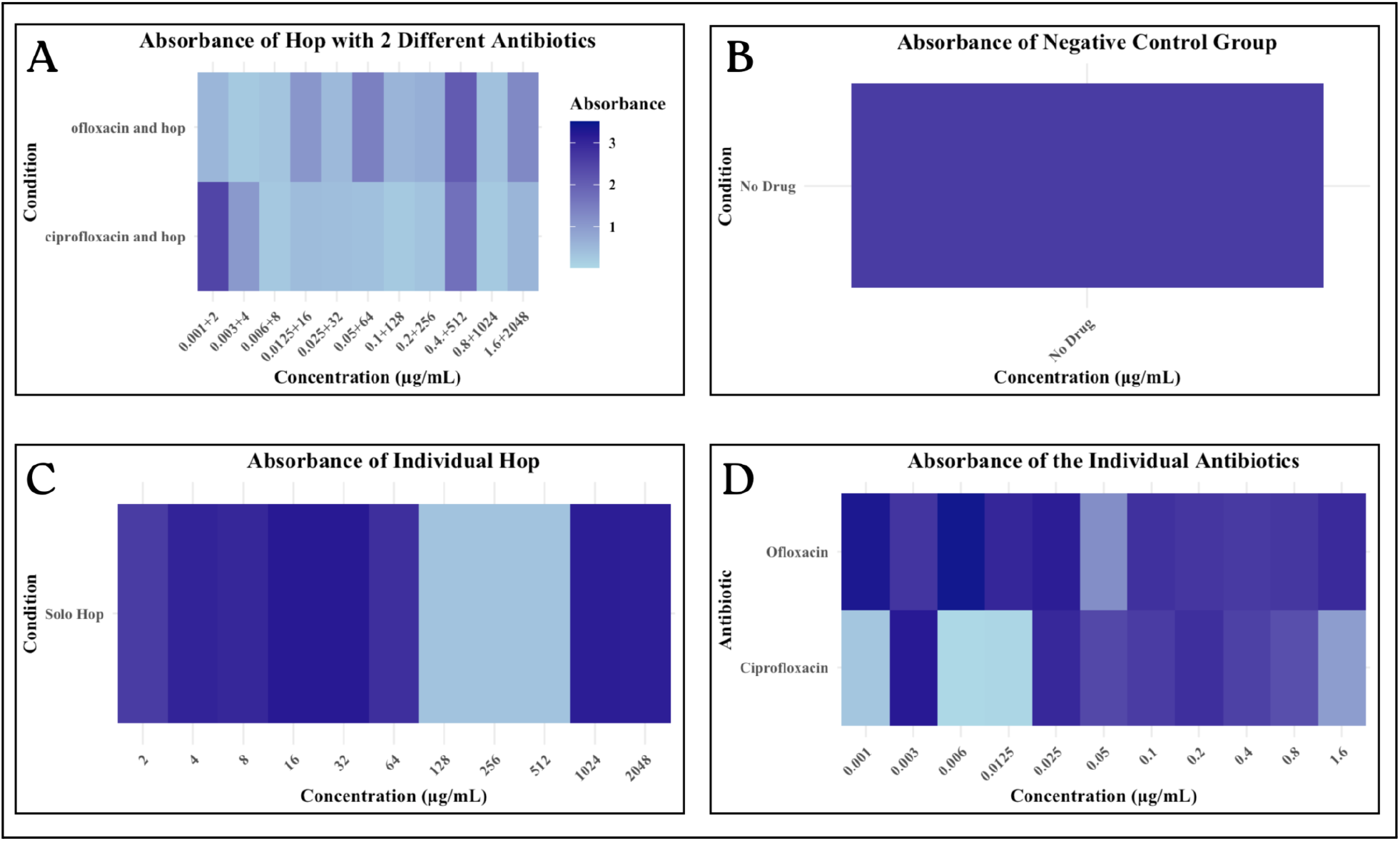
Representation of absorbance levels in CV-stained *S. aureus* cultures grown in the presence of different antimicrobial treatments. Conditions consisted of combination treatments (A), no treatment controls (B), individual Hop treatment (C), and individual antibiotics (D).

## Discussion and Conclusion

This study aimed to determine whether combining QS-inhibiting Hop extract compounds with clinically meaningful ciprofloxacin and ofloxacin would improve antimicrobial activity over using each individually. Here, we found that *S. aureus* biofilms were significantly inhibited compared to treatment with antibiotics alone. This study provides insight into how natural products can potentially mitigate the development of resistance to antibiotics like ciprofloxacin in the highly pathogenic bacterium S. aureus. This study also highlights how adding natural compounds could improve drug effectiveness. This was demonstrated through enhanced activity in combination treatments using Hops+ofloxacin compared to ofloxacin alone. Ofloxacin is currently not an antibiotic used to treat *S. aureus* infections due to its low efficacy in this pathogen [15]. This work highlights potential renewed use of existing antibiotics, potential increased sustainability, and potentially informs improved treatments with better patient outcomes by decreasing AMR. A limitation of this study was that the Hop extract precipitated when in contact with water or media. Because of this, conducting more nuanced and planktonic antimicrobial assays, such as the fractional inhibitory concentration assay (FIC), was not feasible. This limited the ability to quantify additive or synergistic effects for the biofilm inhibition observed in Figure 4. Additionally, due to the precipitation, Hop was diluted using methanol, which tested it as a negative control group to ensure that the antimicrobial effects observed were not due to methanol. As observed through the results, methanol had a very low to no inhibition against *S. aureus*— this suggested that it did not impact the potency of Hop. Current studies on Hop are limited; while the studies mainly focus on oral health, this study concentrates instead on AMR [27]. However, a challenge with using Hop compounds for treatment, especially in intravenous therapy, is the precipitation or cloudiness observed when Hop is dissolved in aqueous solvents. This raises concerns about how such compounds would behave in the bloodstream, potentially leading to harmful precipitates. To address this, an alternative could be Hop extracts in topical treatments or oral formulations, where the precipitation issue would be less of a concern, especially given that the compounds would be absorbed and metabolized differently [28]. *S. aureus* is a significant cause of skin infections, which are often challenging to treat [29]. This combination therapy approach could be promising for treating such infections, avoiding concerns about precipitation in the bloodstream while maintaining antimicrobial effectiveness over time. Unlike studies that typically focus on short-term effects, our study explores the impact of these treatments up to five days post-inoculation, providing a more holistic understanding of their potential. Therefore, our study demonstrates the potential of natural compounds and antibiotics like ofloxacin, which are known to be ineffective against *S. aureus*. Future studies should investigate the bioavailability of these combination treatments through animal models and cytotoxicity assays, and explore these combinations against other resistant bacteria. More studies should focus on leading to effective and sustainable therapies, which would contribute to the global efforts against antimicrobial resistance.

## Authorship Contribution Statement

Sreya Kurup designed and conducted the experiments. She plans to present her data as a poster at the upcoming Annual Biomedical Research Conference for the Minority Scientists 2025 in San Antonio, TX, and wrote the manuscript. Dr. Patrick Taylor assisted in experimental design and reviewed the manuscript. Dr. Barnabe D. Assogba (corresponding author) assisted and contributed to manuscript writing and reviewing. Dr. Paul Adams reviewed the manuscript.

## Declaration of Competing Interest

The authors declare no conflicts of interest related to this manuscript.

## Acknowledgements

The authors sincerely thank Kwantlen Polytechnic University (KPU) for providing a supportive environment that fosters scientific research and collaboration. We are grateful to Ashpreet Kaur, a research assistant in Dr. Assogba’s lab, for her support during the experiments. We also acknowledge Lynsey Baillie’s dedication and efficiency at the Applied Genomics Center (AGC) Lab, particularly her contributions to laboratory management and procuring essential lab items. Her support has positively impacted the outcomes of this study. I am so grateful to Nicole Tumbridge (department Co-Chair) and Dr. Erika Eliason (Associate Dean), who forwarded my email to Drs. Assogba and Adams, making it possible for me to meet and introduce my project to both biomedical research experts at KPU. Finally, we extend our gratitude to everyone who contributed, directly or indirectly, to this research.

## Notes

### Competing Interest Statement

The authors have declared no competing interest.

### Summary of Updates

The abstract is restructured to fit a targeted journal format for review and publication. It is now sectioned into background, objective, methods, results, and conclusions.

### Funding

- Fostering Science Program, UBC, British Columbia, CA
- STEM project grant, British Columbia, CA
- Genome Award, British Columbia, CA

